# Interspecies comparative metagenomics reveals correlated gut microbiome functional capacities among vertebrates

**DOI:** 10.1101/2020.06.15.153320

**Authors:** Christopher A. Gaulke, Courtney R. Armour, Ian R. Humphreys, Laura M. Beaver, Carrie L. Barton, Lucia Carbone, Emily Ho, Robyn L. Tanguay, Yuan Jiang, Thomas Sharpton

**Author notes:** Authors contributed equally to this work (Christopher A. Gaulke, Courtney R. Armour). **Corresponding Author:** Thomas J Sharpton, Department of Microbiology and Department of Statistics, Oregon State University, 97330.

## Abstract

While recent research reveals that the gut microbiome drives vertebrate health, little is known about whether the mechanisms these microbes employ to interact with physiology are consistent across host species. To help close this knowledge gap, we compared gut metagenomes across 10 vertebrate species, including biomedical animal models, to define the inter-species variation in the biochemical pathways encoded by gut microbiota. Doing so revealed gut-enriched pathways conserved across vertebrates, as well as pathways that vary concordantly with host evolutionary history. Overall, the functional capacity of the non-human gut microbiome generally reflects that of humans, though a subset of the pathways encoded by human gut microbiota are not well represented in non-human microbiomes. Collectively, these results support the use of animal models to study the mechanisms through which gut microbes impact human health, but suggest that researchers should cautiously consider which model will optimally represent a specific mechanism of interest.

**Significance:** Efforts to understand how the gut microbiome interacts with human physiology frequently relies on the use of animal models. However, it is generally not understood if the biochemical pathways encoded in gut microbiomes of these different animal models – which define the routes of interaction between gut microbes and their hosts – reflect those found in the human gut. To address this question, we compared gut metagenomes generated 10 different vertebrate lineages. In so doing, our study revealed that non-human gut metagenomes generally encode a set of pathways that are consistent with those found in the human gut. However, some human metagenome pathways are poorly represented in non-human guts, including pathways implicated in disease. Moreover, our analysis identified pathways that appear to be conserved across vertebrates, as well as pathways that are linked to the evolutionary history of their hosts, observations that hold potential to clarify the basis for phylosymbiosis.

## Introduction

Gut microorganisms play a crucial role in promoting health^1^, yet insight into the mechanisms through which gut microbes elicit their beneficial effects remains limited. Animal models have proven to be critical research tools in the effort to uncover and validate specific mechanisms of interaction between gut microbes and physiology^2^. However, it is largely unknown whether the biochemical pathways encoded in the gut microbiome of these animal models are consistent with those encoded in the human gut microbiome, which complicates the translational application of discoveries made using these models. To this end, we conducted a comprehensive comparative metagenomic analysis across 135 individuals spanning 10 vertebrate species, including commonly used animal models. Functional metagenomic profiling identified a core set of 3,368 orthologous protein families that are shared in the gut microbiomes of all examined animals. These shared functions are not simply universally abundant across microbial communities; many are rather enriched in guts relative to free-living communities and possibly represent the functions needed to assemble and maintain a vertebrate gut microbiome. Despite this core functionality, machine learning procedures uncovered phylogenetically congruent patterns of gut microbiome functional diversity, which links vertebrate evolution to the gut microbiome’s functional capacity. Variance-weighted linear models also found that vertebrates generally represent the functional diversity and inter-individual variation encoded in the human gut. However, vertebrate species inconsistently reflected the human gut microbiome at the level of the relative abundance and dispersion of specific microbiome functional modules. Collectively, these results support the use of animal models to study the mechanisms through which gut microbes impact human health, but suggest that researchers should cautiously consider which model will optimally represent a specific mechanism of interest.

## Methods

### Sample Collection

A total of 102 fecal samples were collected from: *Macaca mulatta* (n=10), *Macaca fuscata* (n=6), *Macaca fascicularis* (n=5), *Bos taurus* (n=19), *Canis lupus familiaris* (n=10), *Mus musculus* (C57BL/6; n=16), *Sus scrofa* (n=9), *Rattus norvegicus* (Sprague-Dawley; n=12), and *Danio rerio* (5D; n=15) and stored at −80°C until processing (Supplemental Table 1). Human metagenomes were downloaded from NCBI SRA consisting of 10 Americans (PRJNA48479, randomly selected), 11 Italians (PRJNA278393), and 12 Hadza (PRJNA278393). Environmental samples from ocean (PRJEB1787, surface samples), soil (PRJEB10725), and freshwater (MGRAST mgp5692) were also downloaded. Combined, the data set included a total of 183 samples (Supplemental Table 2).

### Sample processing, library prep, and sequencing

DNA was extracted from fecal samples using the DNeasy PowerSoil DNA extraction kit (Qiagen, Hilden Germany) according the manufacturer’s protocol with an added 10m incubation at 65°C immediately before bead beating to facilitate cell lysis. Samples were homogenized on a vortex genie with a MOBIO bead beater attachment for 20 minutes at full speed. Nextera XT (Illumina, San Diego, CA USA) library preparation was performed by the Center for Genome Research and Biocomputing (CGRB) at Oregon State University. Paired end 150bp sequences were then generated on Illumina HiSeq 3000 (Illumina, San Diego, CA USA). This produced, on average, 2.85×10^7^ +/− 1.27×10^7^ reads for each sample (Supplemental Table 3). Sequencing reads are available through NCBI SRA under project accession PRJNA616045.

### Sample processing and annotation

Each metagenomic library was quality and host filtered using shotcleaner (https://github.com/sharpton/shotcleaner) with default parameters and the closest available host genome (Supplemental Table 1). The resulting high-quality host-filtered reads were trimmed to a maximum length of 100bp to ensure consistency across all samples, and annotated with ShotMAP^3^ using the Kyoto encyclopedia of genes and genomes (KEGG v73.1) database^4^. Details about sequencing depth, quality filtering, and classification rates (Supplemental Figure 1A) are included in Supplemental Table 3.

### Statistical Analysis

ShotMAP^1^ generated abundance data was imported into R and converted to a relative abundance dataframe. Richness, Shannon entropy, and Bray Curtis dissimilarity were quantified with the vegan package. Permutational multivariate analysis of variance (adonis; 5000 permutations) was used to determine if the species labels were associated with beta-diversity of metagenomes. A mantel test was used to quantify the correlation between Bray-Curtis dissimilarity and host evolutionary distance (millions of years) of mammals. To measure individual associations between module abundance and mammalian phylogeny, Pagel’s lambda^5^ was calculated for each module individually.

Indicator species analysis (labdsv) was used to identify modules that are indicators of either host-associated or free-living microbial communities. Indicator species analysis quantifies an indicator value based on the abundance and prevalence of each module^6^; indicator values are close to 1 for a module when it has high abundance and prevalence in the specified group and close to 0 when it has low abundance and prevalence in the specified group. To select which modules indicated host-associated vs free-living communities, we chose functions with an indicator value above 0.6 in one community type and below 0.2 in the other, as well as FDR < 0.05.

We conducted a random forest analysis to assess the ability of the functional diversity of the gut microbiome to classify the community type, species, or order of the samples (randomForest). Module abundance across all samples was used to categorize samples into host-associated or environmental communities, host species, and host order. The out of bag (OOB) error was used to assess classification performance.

To assess the translational relevance of non-human gut metagenomes, we measured how well the gut metagenome’s functional capacity of a non-human lineage correlates with that of humans. Prior work revealed extensive inter-individual variation in the functional capacity of the healthy human gut metagenome^7^, so we reasoned that effective representations the human gut metagenome’s functional capacity should model which protein families are encoded in the gut metagenome, their relative abundances, and their interindividual variation. Accordingly, we developed a weighted regression model that measures the association between these features of a non-human and human gut metagenome. Briefly, we used the multiple metagenomes that were available for each host to compute the mean abundance for each KEGG module across the sampled individuals from a given host. We then weighted each module’s mean relative abundance estimate by the observed interindividual variation for that module in both the non-human and human lineages, which has the effect of adjusting the regression for heteroscedasticity. The corresponding regression slopes produced by this model represent the strength with which the relative abundance and variation of modules observed in a non-human gut metagenome predict the corresponding properties of the modules observed in the human gut metagenome.

Kolmogorov–Smirnov (KS) tests were used to determine how similar the abundance distribution of each KEGG module in each animal model was to the abundance distribution in humans. A significant KS test (p < 0.05) indicates that the two distributions being compared are different, therefore modules were determined to have the same distribution if the *P value* was greater than 0.5.

## Results and Discussion

### Gut microbiome functional capacities are conserved across and phylogenetically correlated with vertebrate species

We observed 15,703 KEGG orthologous protein families (KOs) across the ten vertebrate species and three free-living communities examined. Of these KOs, 2,992 were ubiquitously represented (prevalence = 100%) in all communities, 3,368 in vertebrates, and 3,674 in mammals (Supplemental table 4). Protein family richness (Supplemental Figure 1B) and Shannon entropy (Supplemental Figure 1C) were similar between the communities examined, but richness was elevated in soil and ocean communities while Shannon entropy was consistently higher in zebrafish gut, soil, and freshwater communities (Supplemental Table 5). Functional beta-diversity significantly distinguishes free-living microbial communities from gut microbial communities (Figure 1A; F_1, 182_ = 86.35, R^2^ = 0.32, *P* < 2×10^−4^). Beta-diversity (Supplemental Figure 2) also significantly stratified (F_12, 170_ = 69.13, R^2^ = 0.83, *P* < 2×10^−4^) by community type (e.g. *Homo sapiens*, soil, etc). The association between beta-diversity and community type was not entirely due to the presence of fish and free-living communities as a consistent effect was observed when these community types were removed from analysis (F_8, 111_ = 14.12, R^2^ = 0.50, *P* < 2×10^−4^), suggesting that beta-diversity stratifies mammalian lineages (Supplemental Figure 2). When coupled with contemporary metagenome annotation procedures, the increased sample size per lineage and sequencing depth of the present study affords discovery^3^ of these novel associations as compared to a prior study that resolved a more homogenous distribution of microbiome protein families across mammalian species^8^.

**Figure 1:**
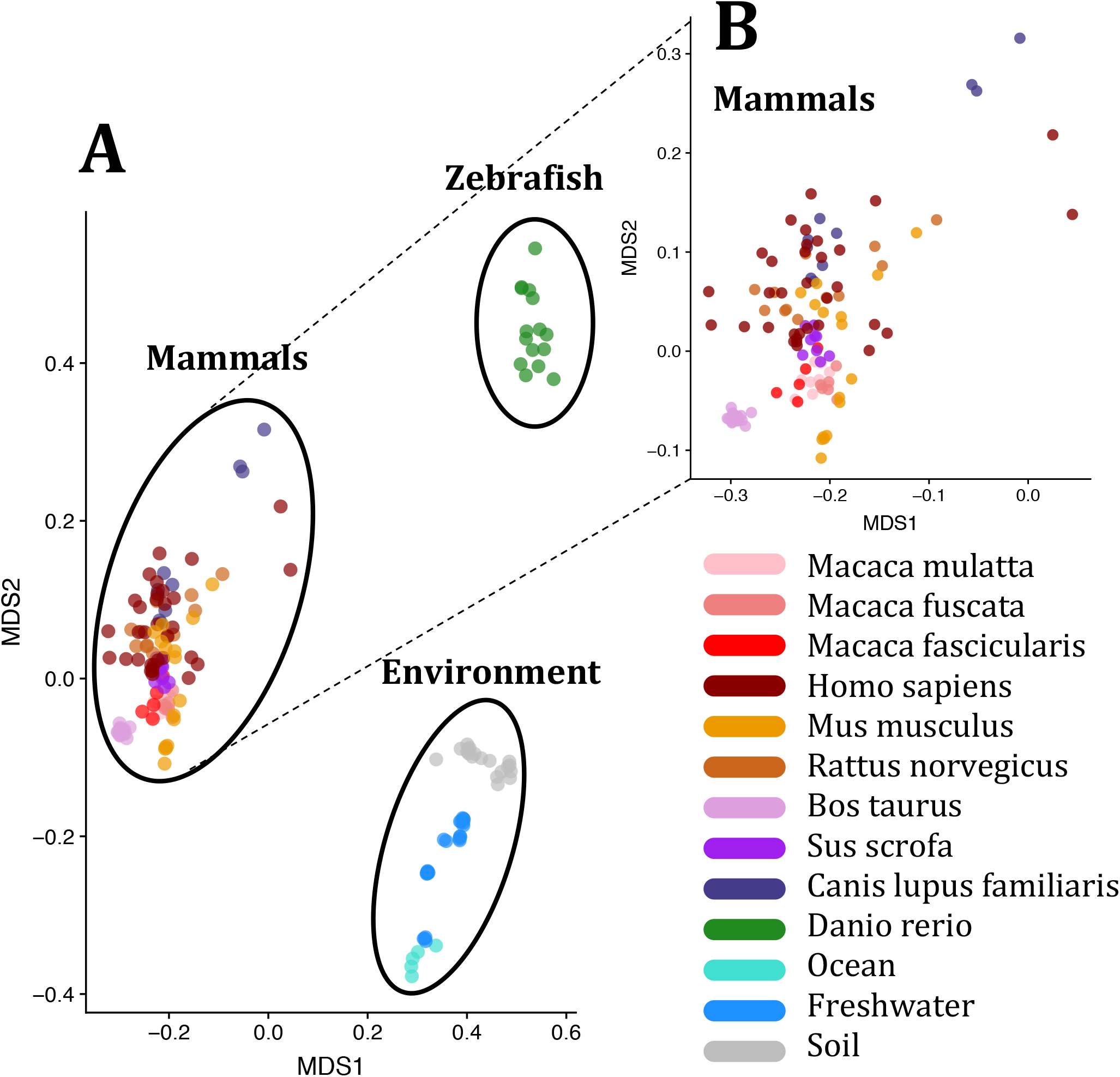
Microbial diversity stratifies mammalian, fish, and free-living microbiomes. **A)** Non-metric multidimensional scaling ordination of host-associated and free-living communities. **B)** Non-metric multidimensional scaling ordination of mammalian gut microbiome diversity. Community type is indicated by point color and community clusters (mammals, fish, free-living) are indicated by ellipses.

Given the association between microbiome functional diversity and mammalian species, we next asked if these patterns were phylogenetically structured. Interestingly, mammalian gut microbiome beta-diversity correlates with host evolutionary distances (r = 0.59, *P* =0.01) indicating that host specific selective pressures, including diet, may shape gut microbiome function (Supplemental Figure 3A). At finer levels of resolution, the abundance of eighty-nine individual KEGG modules associated with mammalian host phylogeny (Supplemental Figure 3B; Supplemental Table 6). For example, phosphotransferase system (PTS), which serve both regulatory and metabolic functions^5^ and have been linked to human disease phenotypes^7^, were significantly enriched in the set of phylogenetically associated modules (odds ratio = 17.26442, *P* = 4.579×10^−10^). Collectively, these results extend previous observations of phylosymbiosis^9^ from taxonomy to functional potential, and suggest that mechanisms such as host filtering^10^, codiversification^9^, and cospeciation^11^ operate to shape the diversification of gut microbiome function in mammals.

### Vertebrates manifest species-specific gut microbiome functional signatures

Given the patterns of beta-diversity stratification (Figure 1) and phylogenetic association (Supplemental Figure 3) between gut microbiome diversity and host species, we next asked if this diversity could distinguish these microbial communities. Random forest classifiers were able to differentiate gut-associated and free-living microbial communities (OOB error = 0%, Supplemental Table 7) as well as individual species (OOB error = 7.4%, Supplemental Table 7). Classification error of individual animal species was primarily driven by an inability to correctly classify macaques at the species level (90% of misclassified samples were macaques, Supplemental Table 7, Supplemental Figure 4). Similar accuracy was observed using other levels of functional annotation as the model feature set (i.e., KO and KEGG pathways; Supplemental Table 7). Since the majority of classification error occurred between closely related lineages, we evaluated if higher levels of host taxonomy yielded a more accurate random forest model. Using host order substantially improved the accuracy of classification (OOB error = 0%, Supplemental Table 7), indicating that distinct lineages in the host phylogeny tend to carry distinct functions in their gut microbiome. We then measured the robustness of this random forest classifier by using it to predict the taxonomy of a gut metagenome’s host at the order level using an independent dataset that includes a diverse array of host lineages^8^. Despite differences in sequencing technology and depth, phylogenetic breadth, and species level taxonomic overlap, our model accurately classified a majority of host lineages in the independent data set (OOB error 31.0%). While our model accurately classified most artiodactyla (67%), carnivora (71%), and primate (82%) samples (Supplemental Table 7), it performed poorly when classifying rodentia samples (OBB error = 100%) possibly due to limited microbial diversity in inbred vs wild rodents^12^. Collectively, these results further suggest that microbiome functional potential is phylogenetically structured.

### Microbiome protein families differentiate gut-associated from free-living communities

Since microbiome functional diversity stratifies free-living and gut-associated communities, we next identified microbial functions that distinguish these community types. We reasoned that functions associated with gut communities may be relevant to survival in the gut or linked with host physiology. We used indicator species analysis to identify KEGG modules that indicate gut-associated vs free-living communities. Of the 658 modules in the full dataset, we identified 65 modules that indicate host-associated communities and 187 modules that indicate free-living communities (Figure 2, Supplemental Table 8). Among the top indicators of free-living communities were modules related to photosynthesis (e.g. M00163/M00161 Photosystem I/II and M00162 Cytochrome b6f complex) and nitrification (e.g. M00528). Gut-associated communities were indicated strongly by phosphotransferase system (PTS, e.g. M00269/M00271) and antimicrobial resistance modules (e.g. M00746/M00738). Over one-third of the gut indicators associate with host phylogeny (Supplemental Figure 3B), suggesting that many microbiome functions enriched in gut-associated communities are subject to ecological or evolutionary forces, such as natural selection, that yield patterns of association with host evolutionary history.

**Figure 2:**
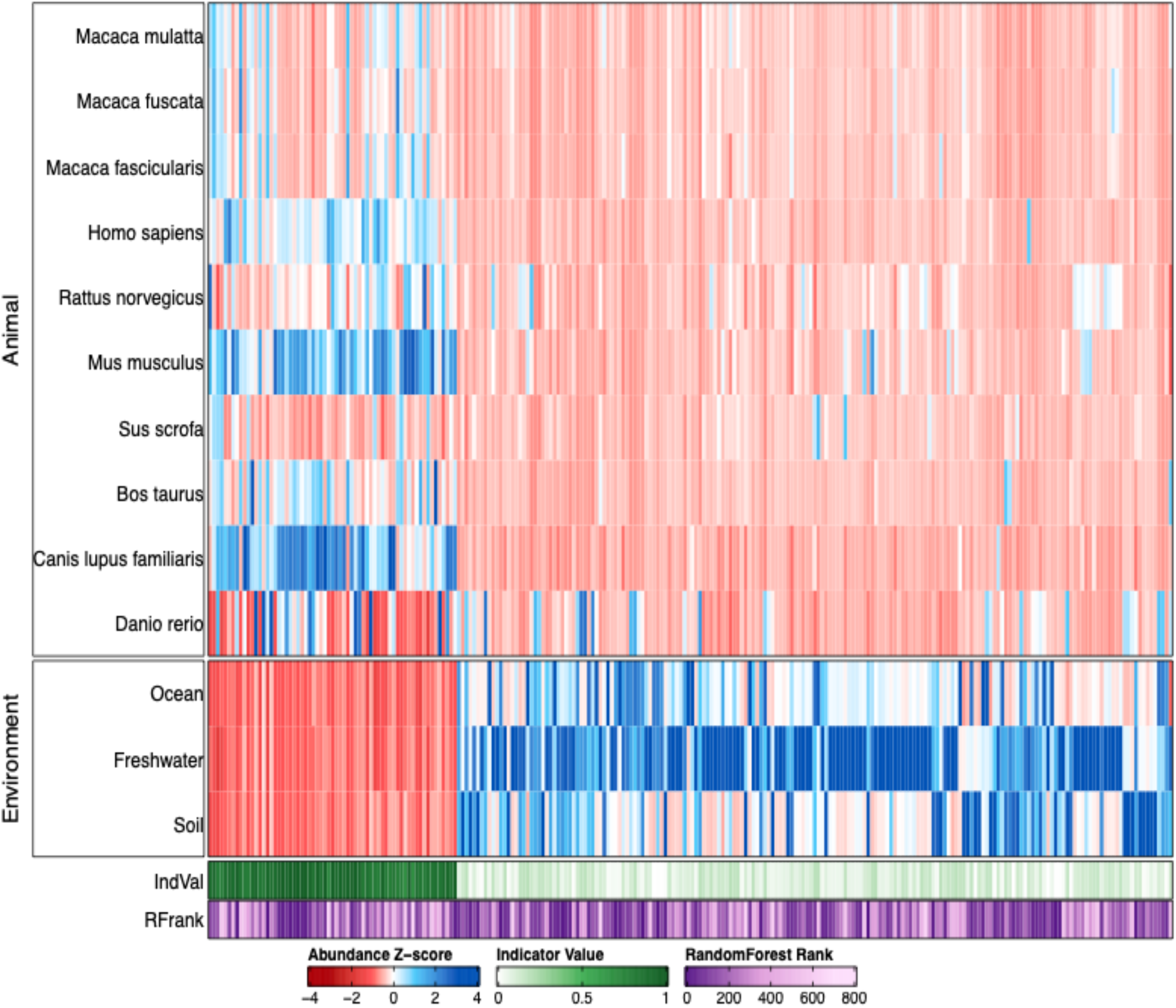
Functional indicators of host-associated or free-living microbial communities. A Heat map of the 252 host-associated or free-living microbiomes indicator modules. Rows are community type, columns are modules, and values are column normalized relative abundance in each community type. The green track is the indicator value for host-associated communities and the purple track is the importance rank from random forests where 1 is most important.

### Recapitulation of the human gut microbiome functional capacity by various vertebrate species

Animal models are widely used to identify interactions between host physiology and the microbiome. However, little is known about the translational relevance of these models to humans. We reasoned that animal model gut microbiomes with functional potential more similar to that of humans were likely to operate in a similar fashion and thus be more translationally relevant. At the level of KEGG modules, linear regression analysis demonstrated that all vertebrate species exhibited moderate-to-strong (R^2^=0.72 − 0.95) associations with human functional potential (Figure 3). The zebrafish microbiome abundance exhibited the weakest association with humans (R^2^=0.72), potentially due to distinct gastrointestinal tract morphology^13^, diet^14^, metacommunity, or phylogenetic distance between humans and zebrafish. However, the association between zebrafish and human module abundances substantially exceeded those observed between free-living communities and humans (R^2^= 0.51−0.64). These results indicate that even evolutionarily divergent vertebrate models, like zebrafish, hold utility as models of human metagenomic functional profiles.

**Figure 3:**
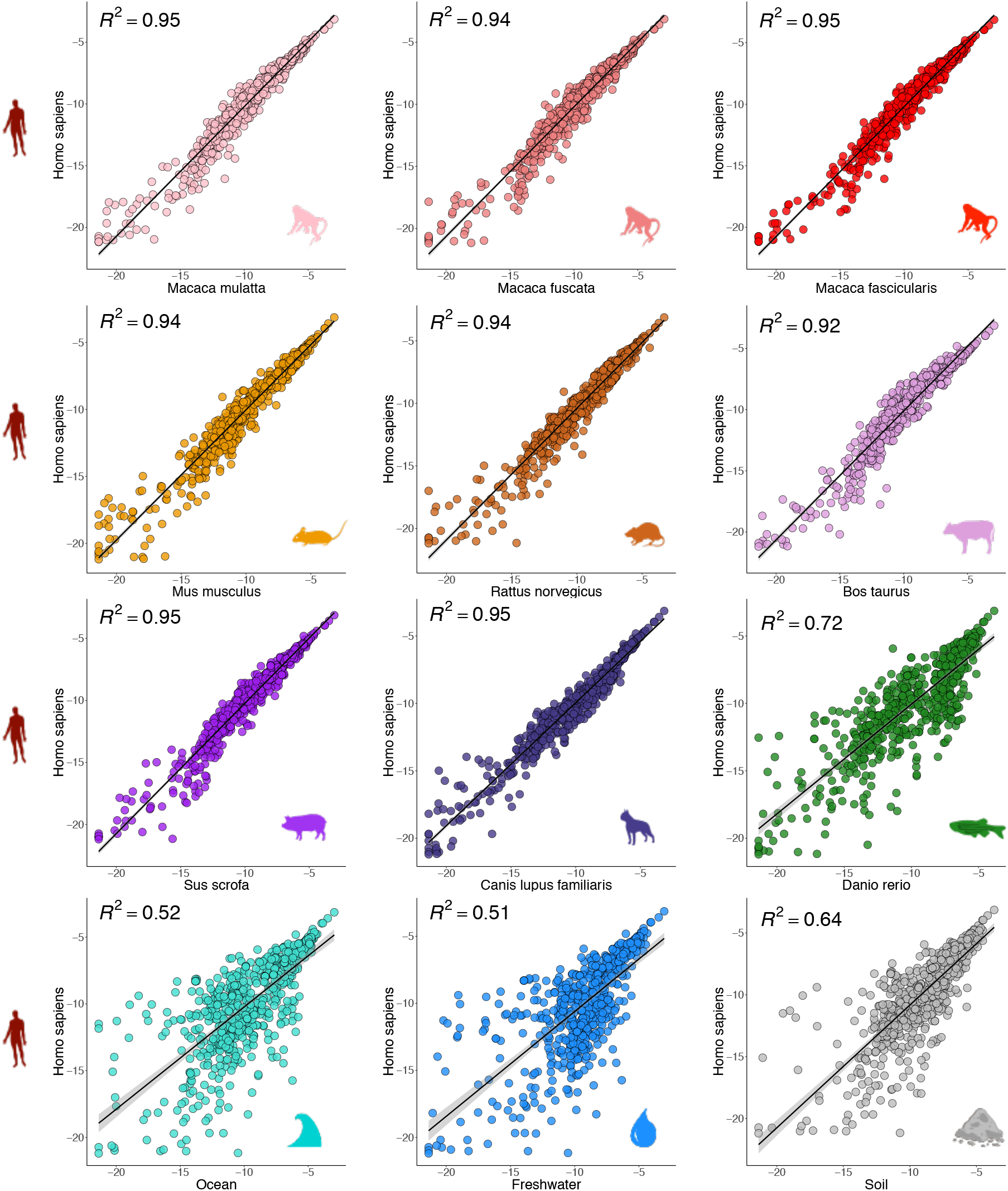
Functional module abundance highly similar between humans and other vertebrates. Scatter plots of human and vertebrate module abundance. Point color indicates the community type that is being compared to humans and R squared of the linear regression is indicated in the upper left corner of each plot. *P value* for all models < 0.05.

**Figure 4:**
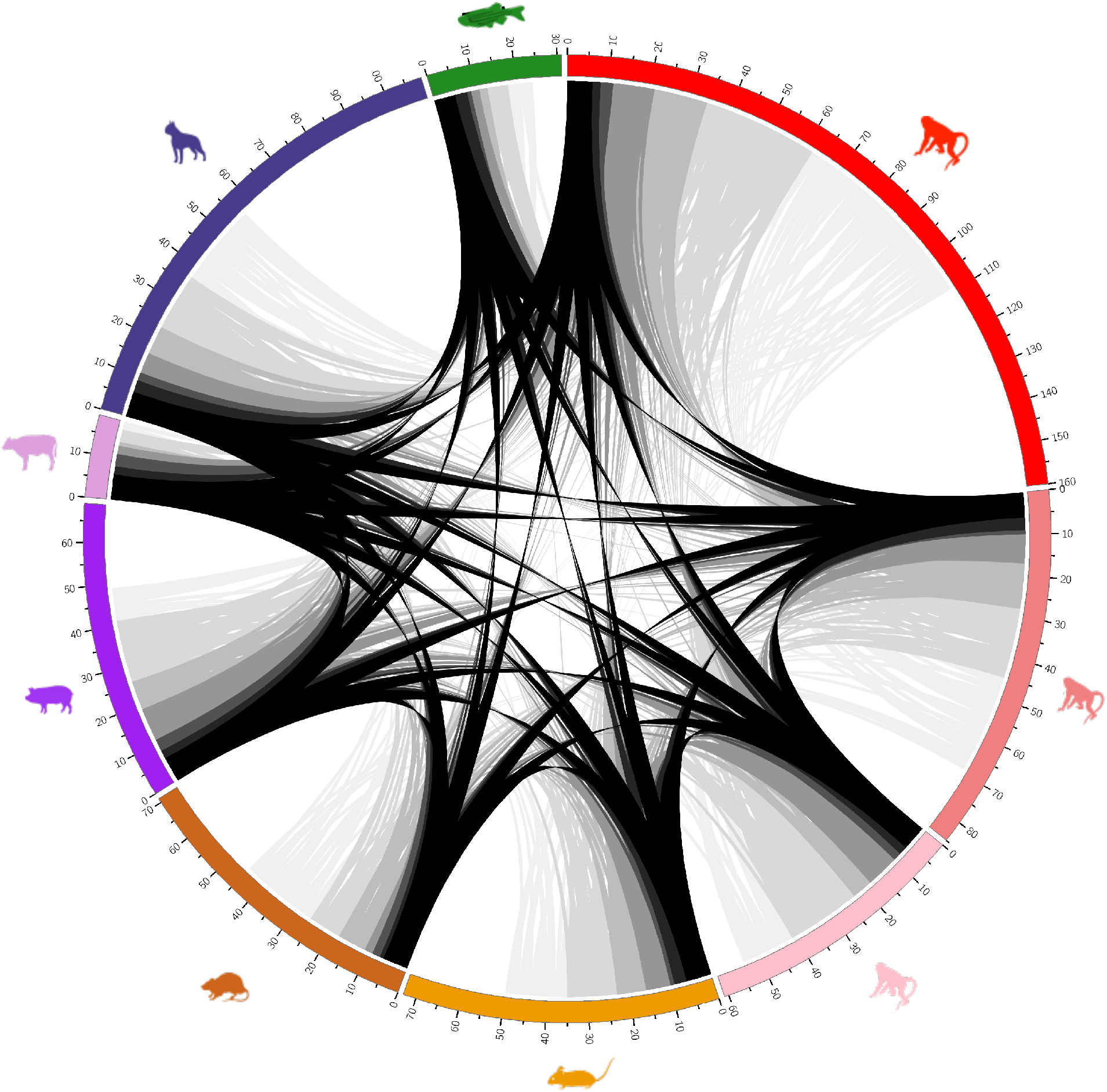
Comparing animal model module abundance distributions to humans. Circos plot of the results from KS tests. The outer track represents the total number of modules for each animal model that has an analogous distribution to humans (*P value* > 0.5). The arcs in the center connect modules for which multiple animal models have an analogous distribution to humans, colored by the number of animal models that share the module. Areas lacking an arc represent modules where that is the only animal model with an analogous distribution to humans.

To more finely resolve the translational relevance of each species, we quantified how well the abundance of each functional module in humans is represented by the different animal models. This information can advance the study of specific human microbiome functions by identifying situations where a particular animal model is most appropriate for the study of specific gut microbiome functions. We conducted Kolmogorov-Smirnov (KS) tests to identify which modules in each non-human host have an analogous abundance distribution to humans (methods, *P* > 0.5). Each of the animal models examined had a set of modules with analogous distributions to that of humans, however the proportion of human modules well modeled by each animal varied. Macaca fascicularis (160) and *Canis lupus familiaris* (109) had the most analogous distributions, while Bos taurus (19) and Danio rerio (31) had the least. Macaca mulatta (61), Macaca fuscata (85), Mus musculus (73), Sus scrofa (69), and Rattus norvegicus (71) all had roughly similar numbers of analogous distributions (Supplemental Table 9). Interestingly, there were several broad functional categories (e.g. antimicrobial resistance, two-component signaling, metal ion transport, and amino acid metabolism) for which no single species was consistently the best model. This result prescribes caution when selecting animal model systems for functional analysis. A total of 154 modules had analogous distributions in multiple species and 202 modules only had one analogous distribution (Supplemental Table 9). Of all human modules, ~46% (302) had no clear analogous abundance distribution amongst other species investigated, indicating that some portion of human microbiome operation may not be modeled well using the animals in our analysis. Included in the modules with no analogous distributions are a number of glycosaminoglycan (GAG) degradation modules (e.g. dermatan sulfate (M00076), chondroitin sulfate (M00077), and keratin sulfate (M00079 and M00080)). Differential abundance of GAG degradation modules in the gut microbiome associated with Crohn’s disease^7^ in humans, however none of the animal models in this analysis appear to consistently represent the human gut microbiome’s distribution of this microbial function. These results demonstrate that consideration should be taken when using animal models to study the gut microbiome in relationship to humans. Not all microbial functions present in the human gut microbiome are well represented by the animal models in this analysis and some are only well represented by specific animals.

## Conclusions

Collectively our study defined the conserved functional diversity of the gut metagenome across commonly used animal models and clarified the translational relevance of these models for studies of host-microbiome interaction. In particular, our analysis uncovered microbial functions that are carried by all examined vertebrates, distinguished host-associated from free-living communities, or that codiversified with host phylogeny. These functions hold great potential to play an important role in gut microbiome operation and possibly interface with host physiology. However, the use of the KEGG database for functional annotation limits our understanding to genes present in the database. While we expect KOs to reflect general trends, many of the genes that cannot be classified into a KO may deviate from the patterns described here. Assembly of metagenomes to create integrated gene catalogues (IGCs) of these samples in conjunction with the existing IGCs^15–20^ could allow the study of novel and understudied microbial functions and their distribution across vertebrates.

Finally, while our results indicated that all evaluated animal models meaningfully estimate human metagenomic diversity, they also prescribe caution when selecting an animal model to study specific mechanisms of host-microbe interactions. The inclusion of more host lineages and greater replication within the included lineages, especially from individuals with specific disease manifestations, will also expand our understanding of model relevance. Ultimately, the selection of the most appropriate animal models should be informed by the metagenomic similarity as well as the physiological relevance and research tools associated with the system.

## Acknowledgements

This material is based upon work supported by the National Science Foundation under Grant No. 1557192, the National Institutes of Allergy and Infectious Diseases of the National Institutes of Health under award number (R21AI135641), the National Institutes of Environmental Health Sciences under award number (R01ES030226), an ACM SIGHPC/INTEL fellowship to CRA, and a Larry W. Martin & Joyce B. O’Neill Endowed Fellowship to CRA. LC is funded by the NIH/OD P51 OD011092 to the Oregon National Primate Research Center. The content is solely the responsibility of the authors and does not necessarily represent the official views of the National Institutes of Health. We thank the Oregon State University (OSU) Swine Facility, the OSU Dairy Farm, the Sinnhuber Aquatic Research Facility, and the Oregon National Primate Research Facility for assistance in provisioning samples.

## Author Contributions

CAG, CRA, LC, EH, RLT, and TJS conceived and designed experiments and analyses. CAG, CRA, LMB, CLB, RLT, IRH, and YJ conducted the experiments and data analysis. CAG, CRA, and TJS wrote the manuscript.

## Supplemental Table Legends

**Supplementary Table 1: Summary of dataset.** Overview of the host species or environments sampled for this analysis. Citations and DOIs are provided for samples acquired from prior publications. Abbreviations as follows:

ONPRC: Oregon National Primate Research Center
OSU: Oregon State University
UCSF: University of California San Francisco
ASPCA: American Society for the Prevention of Cruelty to Animals
SARL: Sinnhuber Aquatic Research Laboratory in Corvallis OR USA

**Supplementary Table 2: Sample metadata.** All available metadata for the samples used in this analysis. Metadata includes sample identifier, the host species (or environment type) of the sample, host order (or environment type), host group (divides samples by their sampling location), host type (animal or environmental), host sex, host breed, sampling location, host diet, and sample accession.

**Supplementary Table 3: Overview of sample processing and annotation.** Summary of the number of starting reads for each sample, how many remained after quality filtering, the sub sampled number, and the classification rate of the subsampled reads to the KEGG database. Some of the larger samples (i.e. soil and ocean) were subsampled prior to processing reduce size footprint and time of processing.

**Supplementary Table 4: KEGG orthology group prevalence across groups.** The prevalence (number of samples with KO/total number in group) of each KO identified in the dataset across groups. A prevalence of 1 indicates some abundance value of the KO in all samples within the group. A prevalence of 0 indicates complete absence of the KO in all samples within the group.

**Supplementary Table 5: Shannon entropy.** Results of pairwise Wilcoxon Rank Sum tests of Shannon entropy between all pairs of sample groups. *P values* are adjusted using the holm correction.

**Supplementary Table 6: Functional modules associated with host phylogeny.** Pagel’s lambda value (Lambda), *P value*, log likelihood (logl), false discovery rate (FDR), and the log-likelihood for lambda = 0, of modules that associate with host phylogeny (FDR < 0.05).

**Supplementary Table 7: Random forest classifier performance across KEGG levels on label types.** A summary of the results from random forest classification of sample type (i.e. animal or environment), species, or order based on KEGG orthologous groups, modules, or pathways. OOB error is the overall error rate for the classifier across all groups and label error is the percent of samples for the respective label that were incorrectly classified.

**Supplementary Table 8: Indicator species analysis summary.** The indicator values of 252 KEGG modules that indicate either animal or environment groups (see methods). KEGG module definition is an abbreviated definition. The “animal” column is the indicator value for animal samples and the “environment” column is the indicator value for environmental samples.

**Supplementary Table 9: Animal models with analogous module abundance distributions to humans.** For each KEGG module identified in the human gut microbiome samples, the animal models (if any) with an analogous abundance distribution for that module based on KS-tests (see methods). NA values indicate no analogous abundance distribution for that module. Host species count is the total number of animal models with an analogous distribution.

---

**Supplemental Figure 1:**
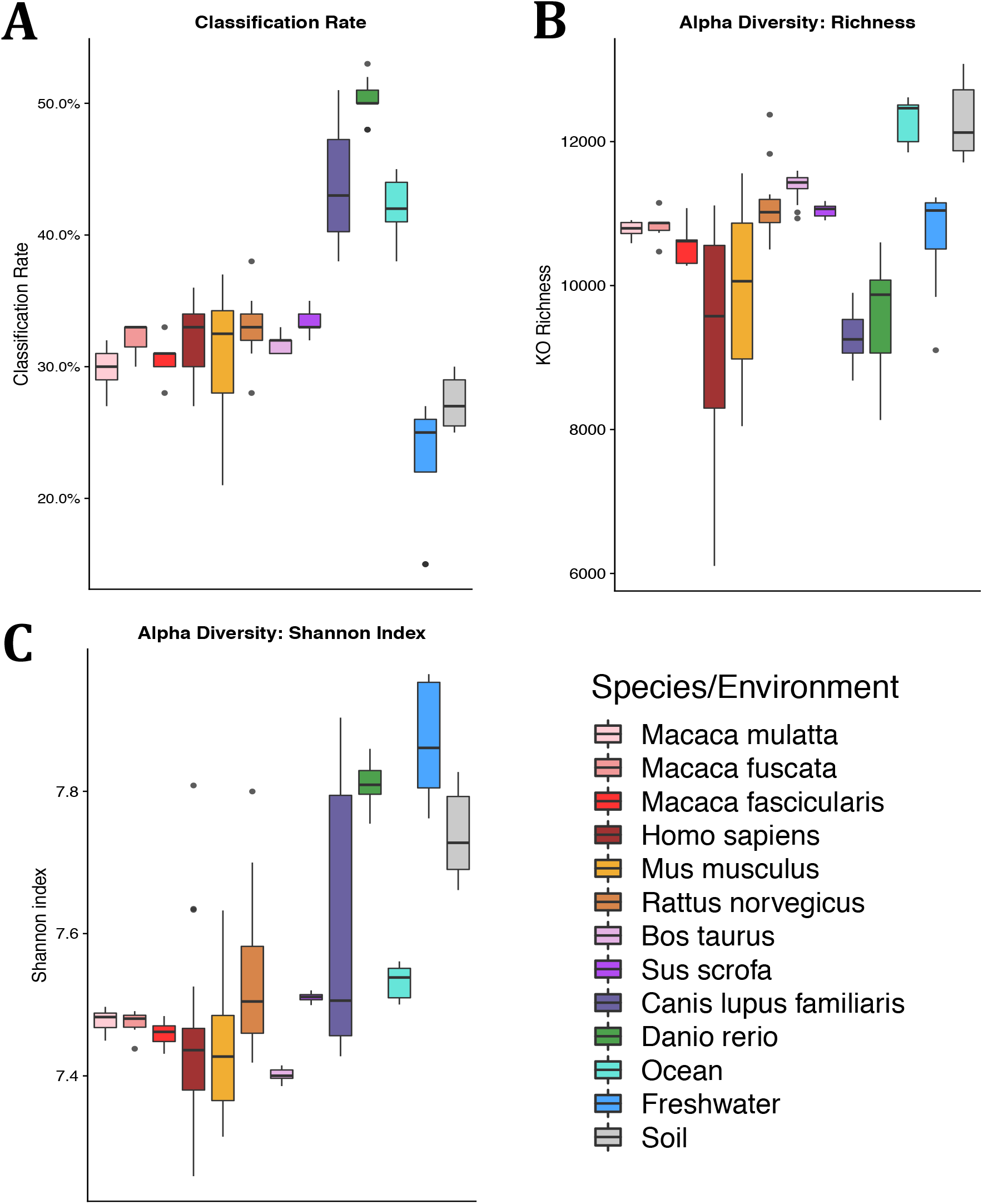
Classification rate and alpha diversity. Boxplots depicting the **A)** Classification rate **B)** richness and **C)** Shannon entropy across the sample groups. Classification rate values for each sample are shown in Supplemental Table 3 and comparisons of Shannon entropy between groups are shown in Supplemental Table 5.

**Supplemental Figure 2:**
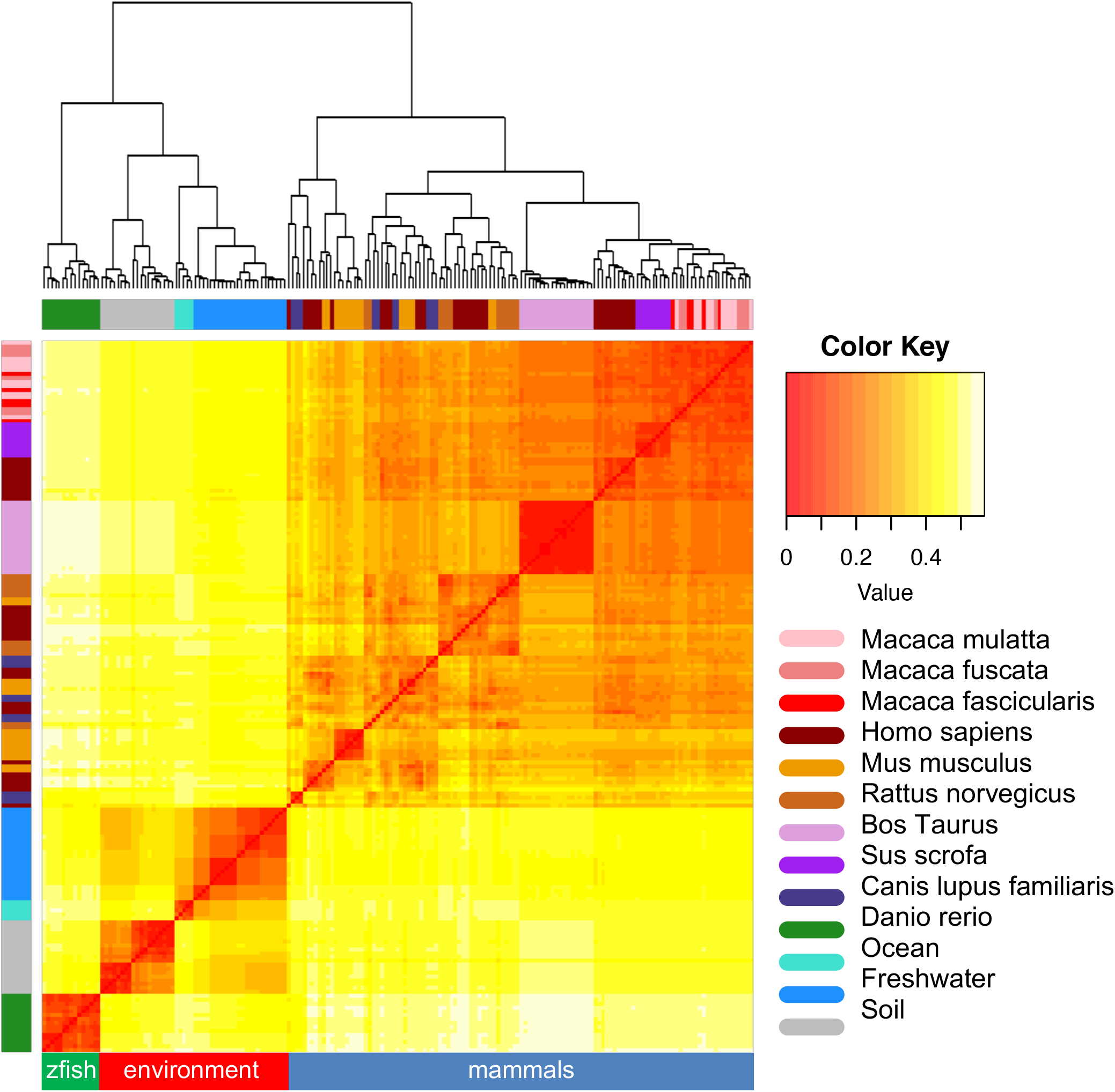
Pairwise beta diversity. Heat map showing the Bray-Curtis dissimilarity between all pairs of samples. A lower value (red) indicates more similarity and a larger value (yellow) indicates more dissimilarity. The samples are clustered by their Bray-Curtis values and structure is depicted by the dendrogram. The y-axis and top x-axis are colored by the sample species or environment label. The bottom y-axis is colored by general sample category of either fish, environment, or mammal.

**Supplemental Figure 3:**
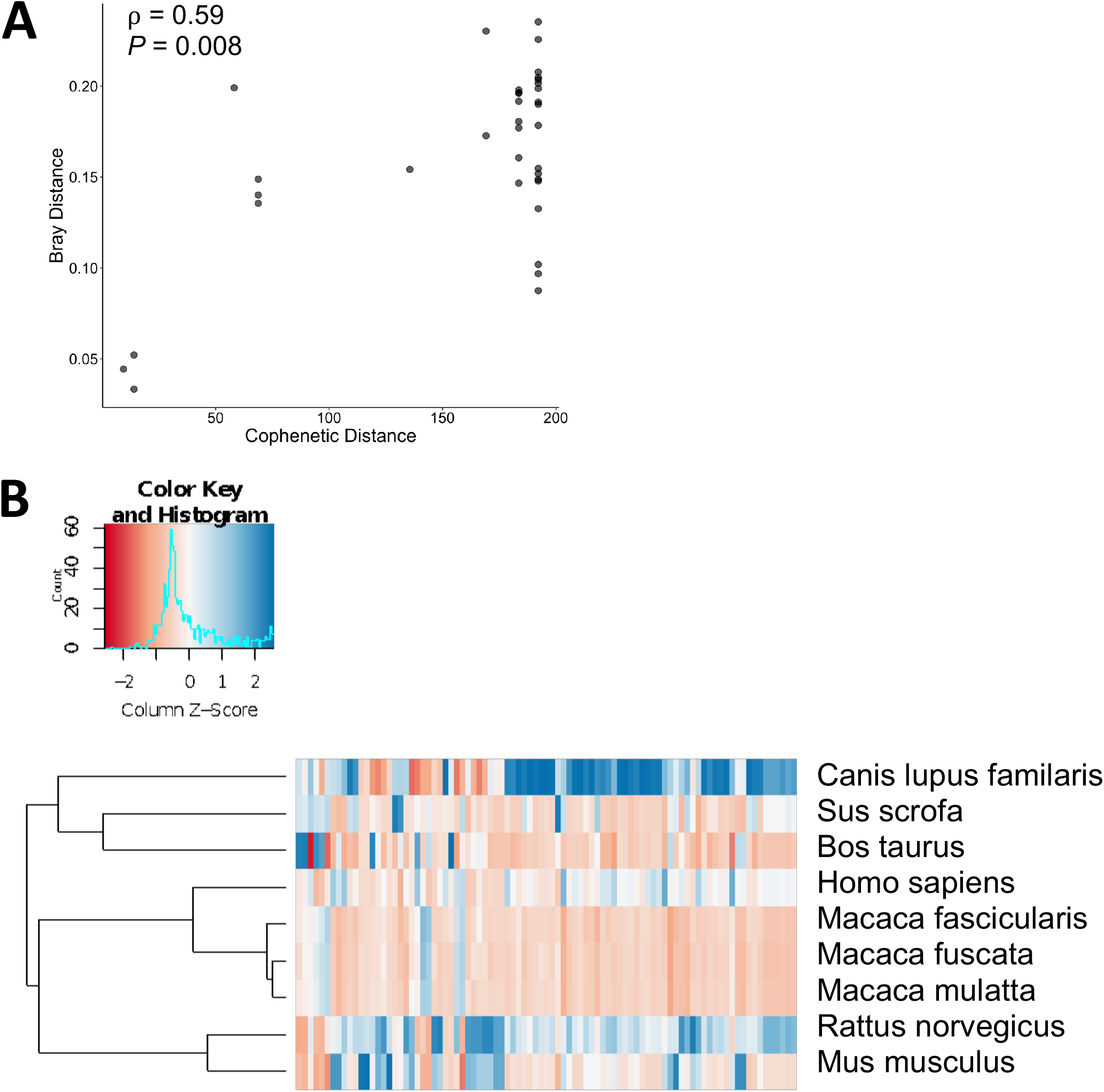
Microbiome functional composition associations with host phylogeny. **A)** Plot comparing distances between microbiome samples based on Bray-Curtis dissimilarity of KO abundance to the phylogenetic distance between hosts. Distances were compared based on a mantel test. **B)** Heat map of scaled module abundance for the modules that associate with host phylogeny based on Pagel’s lambda test.

**Supplemental Figure 4:**
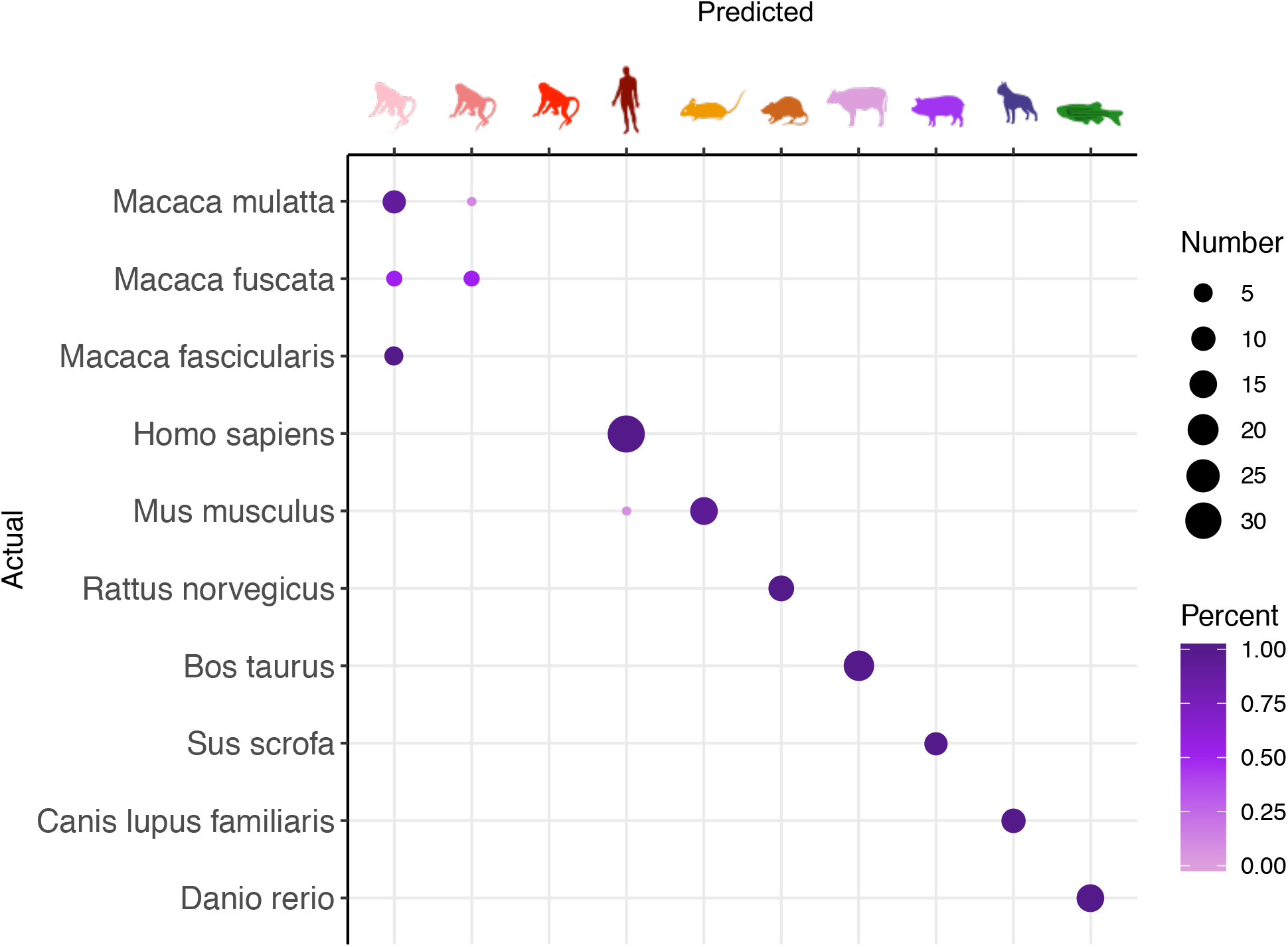
Random Forest. Results from the random forest classifier to classify sample species or environment label based on module abundance. The y-axis shows the sample label and the top x-axis shows the predicted species label (the label ordering is the same on the x- and y-axis). The circles in the plot show how the samples were classified where a dot along the diagonal is represents an accurate classification. Any dots off the diagonal represent incorrect classifications. The size of the dot is representative of the number of samples and the color is the percent of the total samples in the group that number represents.

